# PEA: an integrated R toolkit for plant epitranscriptome analysis

**DOI:** 10.1101/240887

**Authors:** Jingjing Zhai, Jie Song, Qian Cheng, Yunjia Tang, Chuang Ma

## Abstract

**Motivation:** The epitranscriptome, also known as chemical modifications of RNA (CMRs), is a newly discovered layer of gene regulation, the biological importance of which emerged through analysis of only a small fraction of CMRs detected by high-throughput sequencing technologies. Understanding of the epitranscriptome is hampered by the absence of computational tools for the systematic analysis of epitranscriptome sequencing data. In addition, no tools have yet been designed for accurate prediction of CMRs in plants, or to extend epitranscriptome analysis from a fraction of the transcriptome to its entirety.

**Results:** Here, we introduce PEA, an integrated R toolkit to facilitate the analysis of plant epitranscriptome data. The PEA toolkit contains a comprehensive collection of functions required for read mapping, CMR calling, motif scanning and discovery, and gene functional enrichment analysis. PEA also takes advantage of machine learning technologies for transcriptome-scale CMR prediction, with high prediction accuracy, using the Positive Samples Only Learning algorithm, which addresses the two-class classification problem by using only positive samples (CMRs), in the absence of negative samples (non-CMRs). Hence PEA is a versatile epitranscriptome analysis pipeline covering CMR calling, prediction, and annotation, and we describe its application to predict N^6^-methyladenosine (m^6^A) modifications in *Arabidopsis thaliana*. Experimental results demonstrate that the toolkit achieved 71.6% sensitivity and 73.7% specificity, which is superior to existing m^6^A predictors. PEA is potentially broadly applicable to the in-depth study of epitranscriptomics.

**Availability:** PEA is implemented using R and available at https://github.com/cma2015/PEA.

## 1 Introduction

The epitranscriptome, also known as chemical modifications of RNAs (CMRs), is a newly discovered layer of gene regulation with roles in stress responses, RNA folding, and mRNA translation, among other functions (Zhao, et al., 2017). To comprehensively decipher the dynamic epitranscriptome, several high-throughput sequencing (HTS) technologies (**Supplementary Table S1**) have been developed and used to identify CMRs from millions of short reads in particular samples (Helm and Motorin, 2017). However, HTS-based experiments only capture a snapshot of CMRs under certain experimental conditions, and cover just a fraction of the whole transcriptome of a given sample, resulting in the generation of significant numbers of false negatives (non-detected true CMRs) (Luo, et al., 2014). In addition, false positives (wrongly called CMRs) can be induced by the inappropriate use of complex statistical methods and bioinformatics tools. Therefore, integrated tools are critical for the systematic analysis of epitranscriptome data in a standardized, reproducible, and user-friendly manner, particularly those with functionality to accurately identify CMRs at the whole-transcriptome scale.

To our knowledge, RNAModR (https://github.com/mevers/RNAModR) is the only publicly available RNA analytic system specially designed for epitranscriptomics; however, this tool focuses primarily on the location annotation of various CMRs, and lacks functions for CMR calling and prediction. Several CMR predictors have been developed to predict N6-methyladenosine (m6A) and 5-methylcytosine (m5C) (**Supplemental Table S2**); however, the majority was specially designed for analysis of mammalian or yeast sequences, and do not retain their original performance when applied to CMRs from other organisms. This is a particular problem for analysis of CMRs in plant species, because of their lineage-specific sequence and structural properties. These issues underscore the need for integrated tools, particularly for plants, dedicated to general and comprehensive analyses of epitranscriptome data.

In this study, we introduce PEA, a versatile R package for <u>p</u>lants <u>e</u>pitranscriptome <u>a</u>nalysis. Users can easily acquire comprehensive epitranscriptome analytic results, including CMR calling data; transcriptome-scale CMR predictions, generated using an automatically built machine learning (ML)-based classification system; and functional annotation of CMRs. All results can be obtained by step-by-step functions in PEA, rather than requiring users to switch between different analysis tools.

### 2 MODULES AND FUNCTIONS

PEA is composed of three modules: ***CMR Calling***, ***CMR Prediction***, and ***CMR Annotation*** (**Figure 1A**), which can be sequentially executed on multiple processors.

**Fig. 1.**
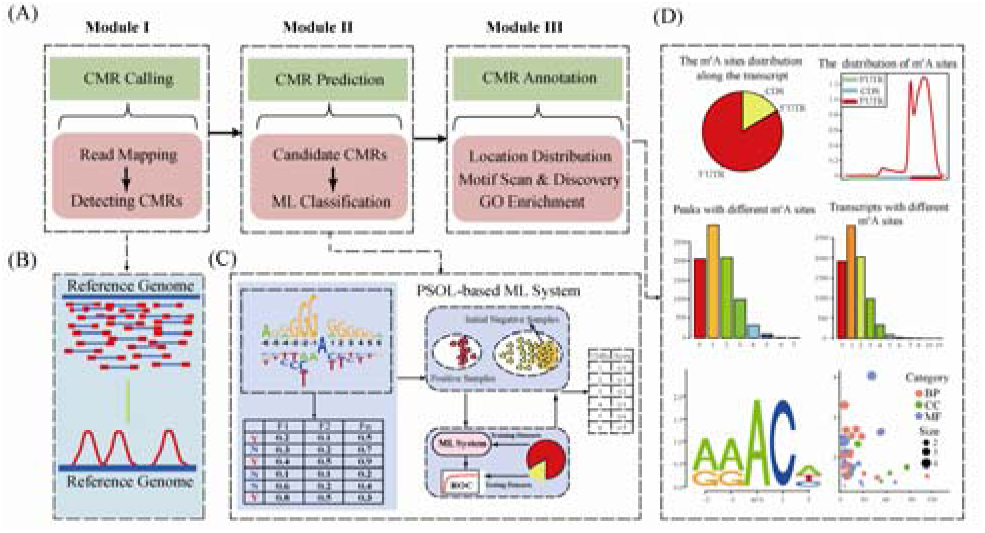
The schematic overview of PEA

***CMR Calling*** (Module I) identifies CMRs from epitranscriptome sequencing data (**Figure 1B**). The input consists of epitranscriptome sequencing reads, and reference genome (or transcript) sequences. The one-step command function “CMRCalling” invokes specific read mapping and CMR calling tools to identify different types of CMRs (**Supplementary Table S3**), in which the quality of read mapping results can be controlled using base-call quality score (e.g., 30 for paired-end reads), and read mapping type (ambiguous or unambiguous). The output consists of identified CMRs in BED format.

***CMR Prediction*** (Module II) predicts CMRs at the transcriptome scale using ML technologies. Considering the sizeable challenge of determining reliable negative samples (non-CMRs), and the mandatory requirement for these for traditional ML classification algorithms, PEA automatically constructs ML-based CMR predictors using PSOL (Positive Samples Only Learning) algorithms, which requires only a set of positive samples (CMRs with high confidence), and iteratively generates a set of reliable negative samples from unlabeled samples (Ma, et al., 2014). A general function, “CMRPrediction”, is provided to implement the PSOL algorithm to automatically build ML-based prediction systems for CMRs of different types and CMRs in different species. Features used to characterize CMRs include **binary encoding**, **k-mer encoding**, and **PseDNC encoding** (see **Supplementary Information**).

***CMR Annotation*** (Module III) is designed to provide insights into spatial and functional associations of CMRs through the function, “CMRAnnotation” (**Figure 1D**). Using this function, the manner of distribution of CMRs in the transcriptome is statistically analyzed, including the spatial distribution of CMRs, and the regions of enrichment of CMRs within transcripts. In addition, motif scanning and *de novo* motif discovery are also provided to investigate the potential regulatory mechanisms leading by CMRs. Moreover, gene functional (Gene Ontology) enrichment analysis is also performed to characterize the enriched functions of CMR-corresponding transcripts using R package “topGO”.

## 3 Application of PEA for Analysis of m^6^A Sequencing Data

To evaluate the efficiency of PEA, we applied it to analysis of *Arabidopsis thaliana* m^6^A sequencing data (see **Supplementary Information**). 11,824 m^6^A peaks were called by PEA from approximately 116 million epitranscriptiome sequencing reads, covering 95.7% (7169/7489) of the peak regions reported in the original paper (Luo, et al., 2014). On this basis, an m^6^A benchmark dataset for training and independent testing was constructed, consisting of 1,674 positive samples (m^6^A modifications) and 36,764 unlabeled samples (m^6^A candidates with RRACH motif). Experimental results from the 10-fold cross-validation on training set demonstrated that the PEA-based m^6^A predictor performed well in identifying m^6^A modifications (AUC = 0.949). A rigorous independent testing showed that the PEA-based m^6^A predictor (sensitivity [Sn] = 71.6%, specificity [Sp] = 73.7%) is superior to recently developed m^6^A predictors for use in plants, such as M6ATH (Chen, et al., 2016) (Sn = 68.0%, Sp = 35.0%) and iRNA-methyl (Chen, et al., 2015) (Sn = 26.5%, Sp = 55.0%). Function analysis of the PEA-based m^6^A prediction at the transcriptome scale revealed the differences in function among genes containing m^6^A in different regions.

## Funding

This work has been supported by the National Natural Science Foundation of China (31570371), the Youth 1000-Talent Program of China, the Hundred Talents Program of Shaanxi Province of China, the Youth Talent Program of State Key Laboratory of Crop Stress Biology for Arid Areas (CSBAAQN2016001), and the Fund of Northwest A&F University.

## Conflict of Interest

none declared.

